# Estimation of Rearrangement Break Rates Across the Genome

**DOI:** 10.1101/402933

**Authors:** Christopher Hann-Soden, Ian Holmes, John W. Taylor

## Abstract

Genomic rearrangements provide an important source of novel functions by recombining genes and motifs throughout and between genomes. However, understanding how rearrangement functions to shape genomes is hard because reconstructing rearrangements is a combinatoric problem which often has many solutions. In lieu of reconstructing the history of rearrangements, we answer the question of where rearrangements are occurring in the genome by remaining agnostic to the types of rearrangement and solving the simpler problem of estimating the rate at which double-strand breaks occur at every site in a genome. We phrase this problem in graph theoretic terms and find that it is a special case of the minimum cover problem for an interval graph. We employ and modify existing algorithms for efficiently solving this problem. We implement this method as a Python program, named BRAG, and use it to estimate the break rates in the genome of the model Ascomycete mold, *Neurospora crassa*. We find evidence that rearrangements are more common in the subtelomeric regions of the chromosomes, which facilitates the evolution of novel genes.

## 1 Introduction

As the molecular revolution allowed scientists to understand the evolutionary import of genetic domains by comparing genes that varied in sequence, the genomic revolution allows us to dissect the importance of genome architecture by comparing genomes that vary in structure.

### 1.1 The Importance of Genome Structure

The chromosomes of both eukaryotes and prokaryotes are known to be highly structured, and that structure is important for gene regulation, the coevolution of genes, and cell division. Moreover, the position of genes within the genome reflects their evolutionary importance. In prokaryotes, genes are biased toward being coded in the leading strand, with conserved essential genes showing higher strand bias (Rocha and Danchin, 2003a,b). Similarly, an emerging trend in eukaryotic genomics has been the concentration of essential genes to the center of chromosomes and the use of subtelomeric regions as sources of genetic innovation (Batada and Hurst, 2007; Brown *et al.*, 2010). A similar trend in the genomics of fungal pathogens is the two-speed genome model, whereby rapidly evolving genes associated with pathogenesis and adaptation to new hosts are found in regions rich with transposons and repetitive elements (Dong *et al.*, 2015).

Given the evident importance of genome structure, it is unsurprising that genomes seem to have evolved mechanisms that stabilize their structure and regulate architectural evolution. In organisms from humans to plasmodium, genes have been shown to be shuffled between the subtelomeric regions, and subtelomeric regions contain unusually high levels of gene duplications and novel genes (Linardopoulou *et al.*, 2005; Cerón-Romero *et al.*, 2018). The presence of transposons in the “high-speed” regions of fungal pathogens have been shown to increase the rate of rearrangements both passively by supplying regions of homology and directly by providing a mechanism for rearrangement (Faino *et al.*, 2016). Rearrangements between subtelomeres are thought to be aided by blocks of repetitive domains, but in humans there is little evidence for direct action of transposons (Linardopoulou *et al.*, 2005).

Within these regions of rapid evolution rearrangement serves as an engine for genetic novelty. In other circumstances, though, rearrangement can act to preserve the genome. Tight linkage of cooperative alleles can ensure co-segregation and co-evolution. Inversions, and perhaps other rearrangements, surrounding cooperative clusters can reduce the risk of separation through recombination as well as prevent repair processes, leading to the development of “supergenes” (Thompson and Jiggins, 2014). Rearrangements are hypothesized to lead to the evolution of specialized functions (Tigano and Friesen, 2016; Brown *et al.*, 2004; Hoffmann and Rieseberg, 2008) and sex chromosomes (Wang *et al.*, 2012; Lahn and Page, 1999; Bachtrog, 2013; Fraser and Heitman, 2004) because of their role in linking advantageous haplotypes. Similarly, recombination is suppressed on the mating-type chromosomes of the fungi *Neurospora tetrasperma* and *Micobotryum lychnidis-dioicae*, and this suppression is linked to an accumulation of rearrangements on these chromosomes (Menkis *et al.*, 2008; Ellison *et al.*, 2011; Badouin *et al.*, 2015). In *N. tetrasperma*, both structural and non-structural suppression have been demonstrated (Jacobson, 2005), although the non-structural mechanisms predate the inversions on this chromosome (Sun *et al.*, 2017).

Under the opposing pressures of maintaining order and generating novelty, the genome structure itself has evolved to promote potentially adaptive rearrangements, and mitigate the harm of maladapative rear-rangements. Similarly, genome defense mechanisms against selfish genetic elements have arisen. For example, in ascomycete fungi Repeat Induced Point Mutation (RIP) acts to silence transposons (Galagan and Selker, 2004). The existence of RIP and structured regions of rapid evolution add to a growing body of evidence indicating that genomes have evolved mechanisms to pick themselves up by their own bootstraps.

Despite the essential functions and evolutionary importance of genome architecture, our understanding of genome architecture has been hampered by the difficulty of assaying the structure of a genome and of inferring the evolutionary history of related genomes. Chromosome mapping techniques such as G-banding, linkage mapping, restriction mapping, happy mapping, and FISH are skill and labor intensive, and only reveal genome organization on the macro- and meso-scales. Genome sequencing can reveal the complete ordering of a genome, but until recently the cost and low quality of sequencing has precluded large scale comparative studies. Now, thanks to the ever decreasing cost and increasing quality of genome sequencing, comprehensive studies of rearrangements of all sizes are possible.

##### Glossary

**affiliations** - *x*^*\**^ - A qbreak is affiliated with a tbreak if the tbreak contains the qbreak. The set of all tbreaks that contain a qbreak are the qbreak’s affiliations.

**breakpoint** - A phosphodiester bond, or set of contiguous bonds, that is present in the reference genome but not the query.

**clique** - A set of vertices in an undirected graph such that each is adjacent to every other.

**clique problem** - The problem of enumerating all maximal cliques in an undirected graph.

**cover** - A set of cliques of a graph whose union includes every vertex in the graph

**history** - *h* - A solution to the tbreak cover problem, which represents a maximum parsimony evolutionary history of the reference genome.

**interval graph** - A graph of a multiset of interval periods, such that each vertex represents an interval over space or time, and intervals that overlap one another are represented by an edge drawn between their corresponding vertices.

**maximal** - A set of elements of a set or graph to that no other element may be added while maintaining a property. With regards to cliques, a maximal clique is a clique that cannot be made larger by adding another vertex and still remain a clique.

**maximum** - A set of elements of a set or graph that is the largest of it’s kind. With regards to cliques, a maximum clique is the largest clique in a graph.

**minimal** - A set of elements of a set or graph that no other element may be removed from while maintaining a property. With regards to covers, a minimal cover is a set of cliques that no clique can be removed from and still cover every element of the graph.

⊂**-minimal** - A qbreak (or vertex of another interval graph) whose affiliations contains no other qbreak’s affiliations as a proper subset.

**minimum** - A set of elements of a set or graph that is the smallest of it’s kind. With regards to covers, a minimum cover is the smallest cover of a graph.

**minimum clique cover problem** - The problem of finding the smallest cover by cliques of a graph.

**multiset** - {*a, b, b*} - An unordered collection of elements which may be, but are not necessarily, unique.

**powerset** - *𝒫* (*X*) - The set of all subsets of a set.

**qbreak** - A region of the reference genome between two orthologous segments that are adjacent in the reference but not in the query.

**set** - {*a, b, c*} - An unordered collection of unique elements.

**tbreak** - *τ* - A clique of qbreaks that is also tree consistent.

**tree consistent** - A tbreak is tree consistent if it contains all and only qbreaks from queries that lie on one side of one branch in the phylogenetic tree.

### 1.2 Phylogenetic Inference of Rearrangements

A dizzying array of new high-throughput assays are enabling the study of genome architecture, but analysis of these piles of data has become the primary challenge. Measuring the pace of rearrangement within different chromosomal domains would ideally involve reconstructing the history of those domains and counting the occurrences of different rearrangements. However, the difficulty of inferring rearrangements makes this task impractical for the increasingly large datasets now available.

Phylogenetic inference is dependent upon the inference of homology between the sequences under study, that is, the alignment. Sequence alignments represent sequences as rows in a matrix where columns represent a homology relation between characters in the sequence. Unfortunately, while pairwise alignment is relatively efficient, the problem of multiple sequence alignment (MSA) is known to be NP-hard (Elias, 2006) and remains a computational challenge. Even matrix sequence alignments, though, are insufficient for aligning genomes because they cannot account for rearrangements between the sequences.

One approach to multiple genome alignment (MGA) is to find and align collinear regions between the genomes, then represent each genome as an ordering on an improper subset of the collinear regions. Inferring rearrangements from the ordering then requires combinatorial analysis. Graph based MGAs improve this paradigm by representing collinear segments as nodes and each genome as a path between the nodes. Edges in such a graph thus represent adjacencies between sequences, or nonhomologous phosphodiester bonds. MGA graphs have the elegant property that rearrangements are equivalent to swapping edges in the graph, and reconstructing possible histories of rearrangements is thus equivalent to converting this multigraph into a graph where all genomes follow the same path (Alekseyev and Pevzner, 2009). Reconstructing genome histories in this way is known as the Multiple Genome Rearrangement Problem (MGRP), which is known to be NP-complete in at least some cases (Caprara, 1999), and the complexity of MGRP balloons as more genomes are added since each genomr adds both more observed rearrangements that fragment the collinear regions and another ordering of those regions.

Furthermore, this sort of MGA suffers from the philosophical problem of splitting MGA into two steps with different rules: first the NP-hard MSA that assumes no rearrangement, and second the NP-complete analysis of rearrangements that assumes the truth of the MSAs. This paradigm conflicts with the biological truth of evolution, which operates under a single set of rules. While we classify mutations (e.g. as single nucleotide polymorphisms, deletions, or transpositions) based upon our observation of the products of evolution (i.e. sequences), these classifications have an ambiguous relationship to the mechanisms that caused them. Cactus graphs, a recursive alignment graph where each node is itself an alignment graph, allow for iterative refinement of the inferences of homology in the context of the larger genome structure (Paten *et al.*, 2011). Cactus graphs thus provide a unified model of homology that accounts for both changes in the nitrogenous bases of the DNA and the phosphodiester bonds that link them.

### 1.3 Estimating Rearrangement Rates

Even given a perfect alignment methodology, there are often multiple equally good solutions to the MGR problem, leading to significant ambiguity in the sequence of events (Sankoff and Blanchette, 1998).

Distance based phylogenetic methods can provide an imperfect solution by counting the number of double-strand breaks between two genomes. Breaks occur when the phosphodiester backbone of both strands of a chromosome are broken and not reformed. When chromosome number is maintained, breaks only become permanent when two double-strands are broken and the ends of the DNA fragments are swapped. Thus, the break rate is less than or equal to twice the rearrangement rate (Sankoff and Blanchette, 1998). Every missing adjacency in a comparison of genome orderings or every branch point in an MGA graph represents a **breakpoint**, and breaks are thus easily quantified.

Distance based methods, though, become less reliable at larger phylogenetic distances. While the risk of reversion of rearrangements is lower than with substitution mutations, alignments between distantly related genomes are more difficult to achieve and they become more gappy, which can lead to an underestimate of break distances. Furthermore, information about the local density of breaks is lost when reduced to a simple count.

Rather than attempt to infer and quantify the number of rearrangement events along a phylogeny, we re-frame the problem to measuring the degree of conservation, or conversely fragility, of the set of phosphodiester bonds within a genome. Rearrangements by their nature break one set of bonds and form a new set of bonds, and so rearrangements represent a decay process on the set of phosphodiester bonds. The fragility of bonds in this context is underlain by the chemical instability of the bonds, but perhaps more importantly reflects selection that acts to carry those changes into the next generation or eliminate them from the population.

Within this framework, we present a method to estimate the Break Rates Across a Genome (BRAG). BRAG uses pairwise alignments between the genome of interest, termed the reference, and a set of related genomes, thus obviating the need for a difficult multiple genome alignment, or even all-against-all pairwise alignment. BRAG employs a novel, interval graph based approach for which efficient algorithms have been previously described. Additionally, BRAG provides a detailed survey of break rates across the genome by computing the likelihood landscape of the break rate at every site in the genome.

## 2 Methods

Here, we first describe the BRAG method in detail. We then employ BRAG to examine the pattern of rearrangements in the genome of the model filamentous ascomycete mold, *Neurospora crassa*. *Neurospora*’s relatively small haploid genomes with low repetitive content, as well as ease of sampling diverse species, make it an attractive model for genomics and the study of evolution due to the ability to easily obtain and sequence many genomes (Gladieux *et al.*, prep; Palma-Guerrero *et al.*, 2013; Heller *et al.*, 2016; Stajich *et al.*, 2009). The extensive knowledge of *N. crassa* biology also allows us to correlate the break rate to other features of the genome with the aim of dissecting which factors affect rearrangement dynamics.

### 2.1 Genomes and Phylogeny

We utilized a dataset of 15 *Neurospora* genomes and one *Sordaria* genome (Nowrousian *et al.*, 2010) to serve as the outgroup. Sequencing and assembly methodologies, assembly statistics, and references are described in Table S1. All new genomes in this study were sequenced using 150bp paired end reads at the Vincent J. Coates Genomics Sequencing Laboratory at UC Berkeley and assembled using the A5-miseq pipeline (Tritt *et al.*, 2012; Coil *et al.*, 2015), following the methodology laid out in our companion paper (Hann-Soden *et al.*, prep). We selected the well studied and complete genome of *N. crassa* (Galagan *et al.*, 2003) as the reference genome.

Since the BRAG methodology attempts to place break events on the branches of a phylogenetic tree to maximize parsimony, using the correct tree topology is essential to accuracy. Furthermore, the break rates estimated by BRAG are scaled by the branch lengths of the tree, so results are biased by inaccurate branch lengths. An accurate alignment and high quality tree are therefore essential to correct inference.

We identified orthologous sequences between the reference and all queries in a pairwise manner with the MUMmer package (Kurtz, 1999; Delcher *et al.*, 1999, 2002; Kurtz *et al.*, 2004), first finding maximal alignments with nucmer with a gap length of 2000bp, then filtering for the highest scoring 1-to-1 set of alignments with delta-filter. We filtered for regions of the reference that had aligned orthologous regions to all queries. We then performed multiple sequence alignment between orthologues of all such regions using MAFFT (Katoh *et al.*, 2002; Katoh and Standley, 2013) using the –globalpair setting for global alignment, 1000 maximum refinement iterations, and the JTT 10 model (Jones *et al.*, 1992). We discarded regions on the ends of alignments for which any sequence had missing data before concatenating all alignments into a single 11,400,437 character alignment with 1,648,053 parsimony-informative sites. We partitioned the concatenated alignment into bins by their approximate rate of evolution using TIGER (Cummins and McInerney, 2011), which we re-wrote to fix errors, optimize for genome-scale alignments, and bin using the method described by Rota *et al.* (2017). Finally, we searched for the maximum-likelihood tree with IQTree using the -m MFP option for model selection, generating 1000 ultrafast bootstraps, and estimating branch support using 1000 replicates in the non-parametric Shimodaira-Hasegawa-like approximate Likelihood Ratio Test (SH-aLRT) (Nguyen *et al.*, 2015; Kalyaanamoorthy *et al.*, 2017; Thi Hoang *et al.*, 2017; Chernomor *et al.*, 2016).

We provide code for the phylogenetic analysis and our rewrite of TIGER in the Data Access section of this paper.

Our whole genome phylogeny of *Neurospora* is strongly supported by both metrics (Fig. 1), and thus should be suitable for BRAG analysis.

**Figure 1:**
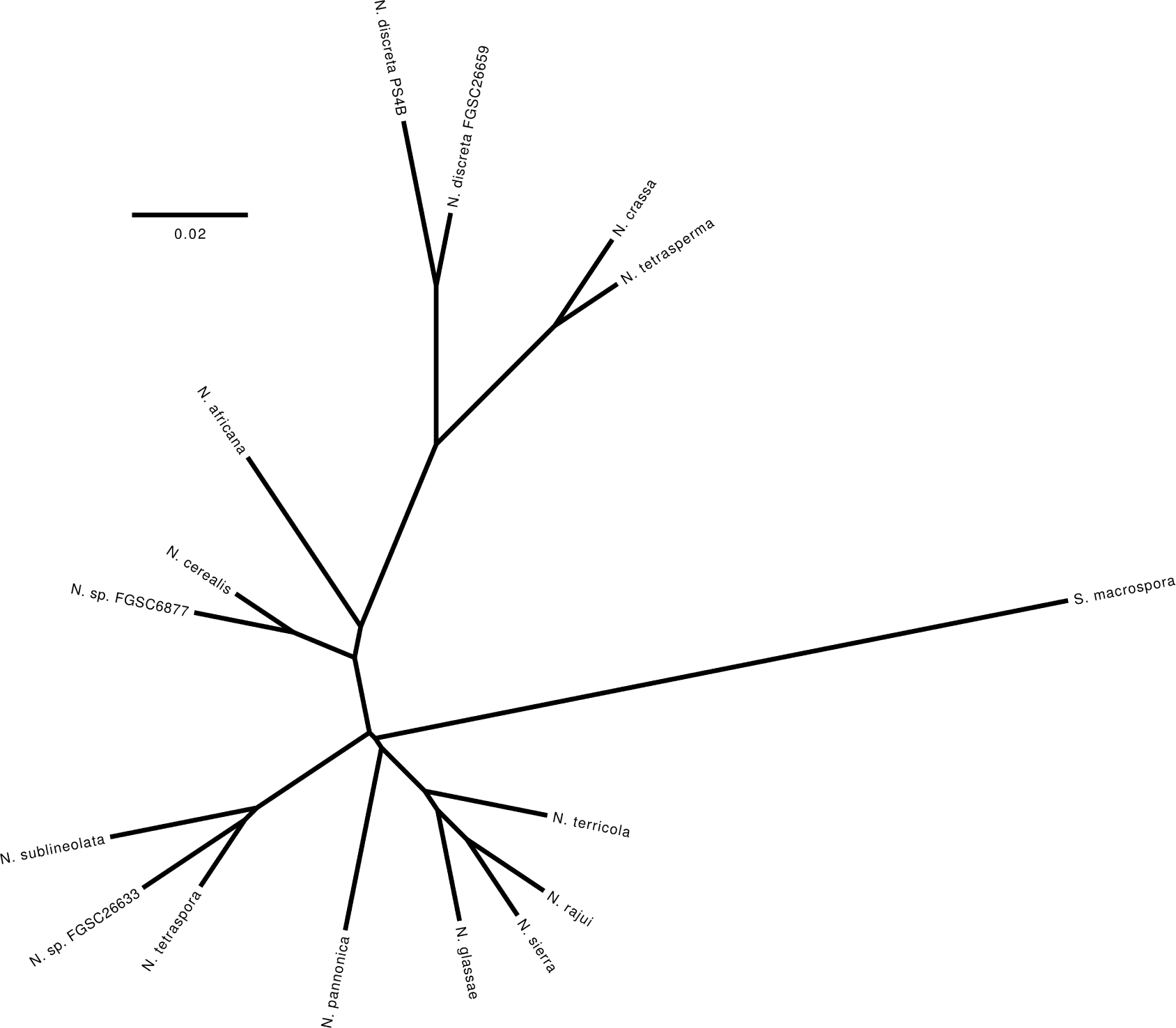
The whole genome phylogeny of all *Neurospora* species and an out-group (*Sordaria macrospora*) for which at least one genome is available. All branches have ¿98% bootstrap and 100% SH-aLRT support.

### 2.2 Whole Genome Alignment

For BRAG analysis, we roughly aligned all the genomes to the *N. crassa* reference genome using Murasaki and OSfinder (Popendorf *et al.*, 2010; Hachiya *et al.*, 2009). Murasaki identifies roughly similar short sequences, or anchors, between genomes. We configured Murasaki to find 36-mer anchors with at least 28 matches. OSfinder identifies collinear chains of anchors and attempts to find the most likely set of orthologous segments from non-overlapping chains. We configured OSfinder to find orthologous segments at least 1,000 bp long.

#### 2.2.1 Query Breaks

For each pairwise genome alignment we define breakpoints to a query (or **qbreaks**) as the regions on the reference genome between two orthologous segments (Fig. 2). Qbreaks have both left and right endpoints and a query, with the property that no two qbreaks of the same query are overlapping.

**Figure 2:**
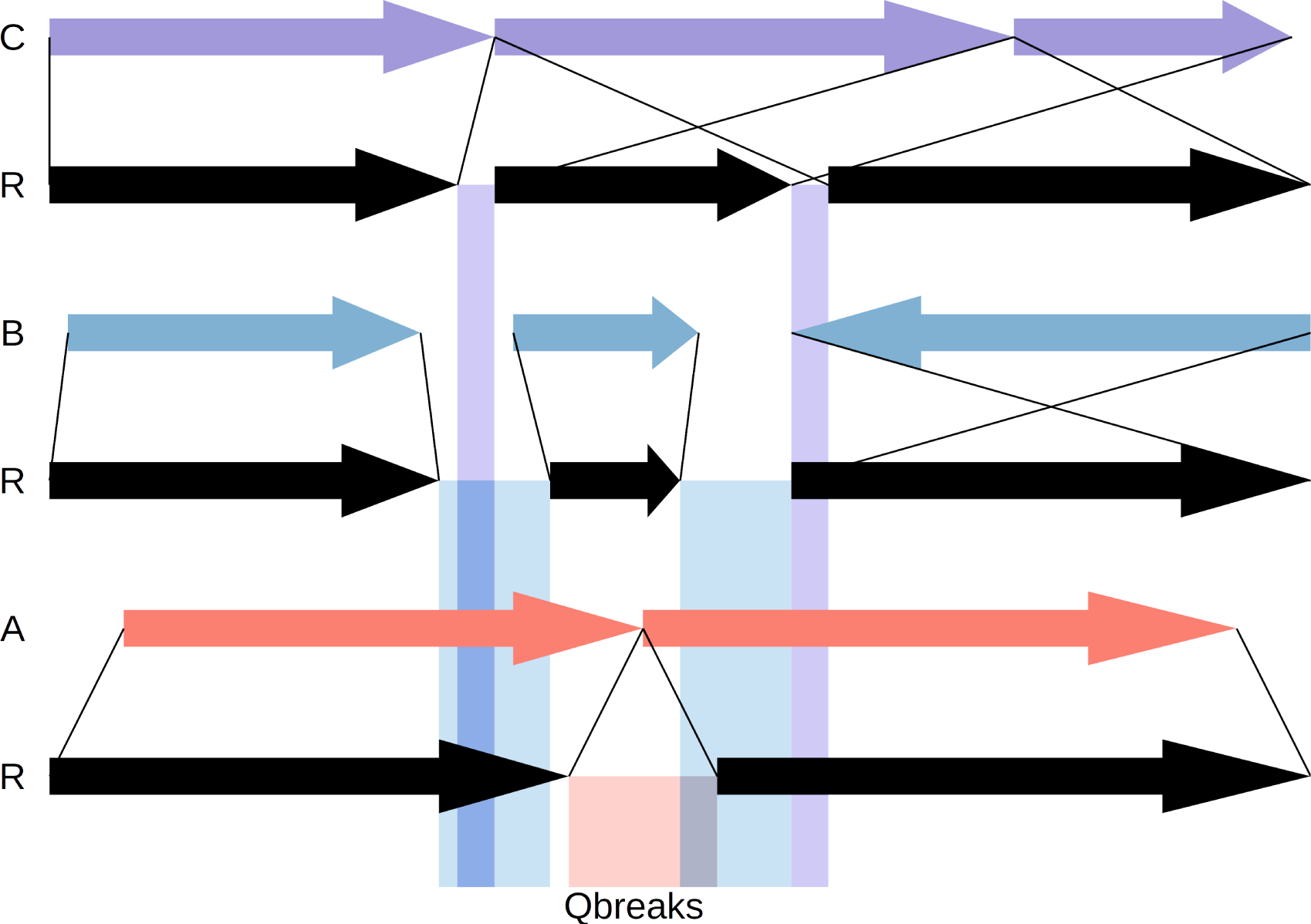
Three toy alignments between query genomes (A, B, and C) and the reference (R). Orthologous segments between the reference and each query are shown as colored block arrows, with thin black lines showing the orthology relationship. Qbreaks are represented as shaded rectangles emanating from the reference at regions between orthologous segments, and colored by their corresponding query alignment. No two qbreaks of the same color can overlap, but qbreaks of different colors (i.e. queries) can overlap.

Orthologous segments are defined by sequence, or nucleotide base, identity and are closed intervals on the reference sequence. As such, breakpoints occur on the bonds between nucleotides and qbreaks are the open intervals between the orthologous segments. When two orthologous segments are immediately adjacent there is thus a qbreak between them encompassing a single bond.

Additionally, due to fragmentation of the query genomes, orthologous segments that are on the ends of the scaffolds could be adjacent on a chromosome or not. We define qbreaks where one or both orthologous segments are internal on a scaffold, and thus their adjacencies are known, as “true”. Conversely, qbreaks where both orthologous segments are on the ends of a scaffold in the query as “false”, equivalent to “can’t tell” vertices described by Zheng and Sankoff (2016). As a conservative estimate, we calculate break rates utilizing only “true” qbreaks, which represents a lower-bound on the true break rate. For an upper-bound estimate, we calculate the break rate using both “true” and “false” qbreaks.

### 2.3 Identification of Breakpoints

Given the set of qbreaks, we seek to identify double-strand breaks that have occurred along the branches of the evolutionary tree connecting the reference to the queries. Multiple qbreaks can be explained by the same double-strand break event when they overlap and when their queries share a branch leading to the reference. Under the paradigm of maximum parsimony, we favor these breaks that can explain the existence of the most qbreaks with the fewest number of events. Similarly, we do not consider the possibility of reversions, which would require the same sites to break again at the same time and the ends of the fragments to exchange back to their original orientation and adjacencies - an event we assume to have a vanishingly small probability.

To this end, we frame the problem in graph theoretic terms and find that it is a case of the **minimum clique cover problem**. Well known in mathematics as one of Karp’s original 21 NP-complete problems (Karp, 1972), the minimum clique cover problem is any problem that is equivalent to the following: Knowing which students at a school share a class, find the minimum number of classes that must be offered and which students are in which classes.

#### 2.3.1 Tree Consistent Breaks

We define a tree consistent breakpoint (or **tbreak**) as a set of overlapping qbreaks whose queries form a **tree consistent** partition (Fig. 3). In the manner of Alekseyev and Pevzner (2009), a partition of queries is tree consistent if it or its complement form a monophyletic group. That is, a partition of queries is tree consistent if it is a bipartition present in the tree. A tbreak’s placement in the tree corresponds to this bipartition (or branch). We use *τ ∈* T to denote a tbreak within the set of all tbreaks. Since tbreaks are bounded by their member qbreaks, the left endpoint of a tbreak (*min*(*τ*)) is the maximum left endpoint of it’s members. Analogously, a tbreak’s right endpoint (*max*(*τ*)) is the minimum right endpoint of it’s members.

**Figure 3:**
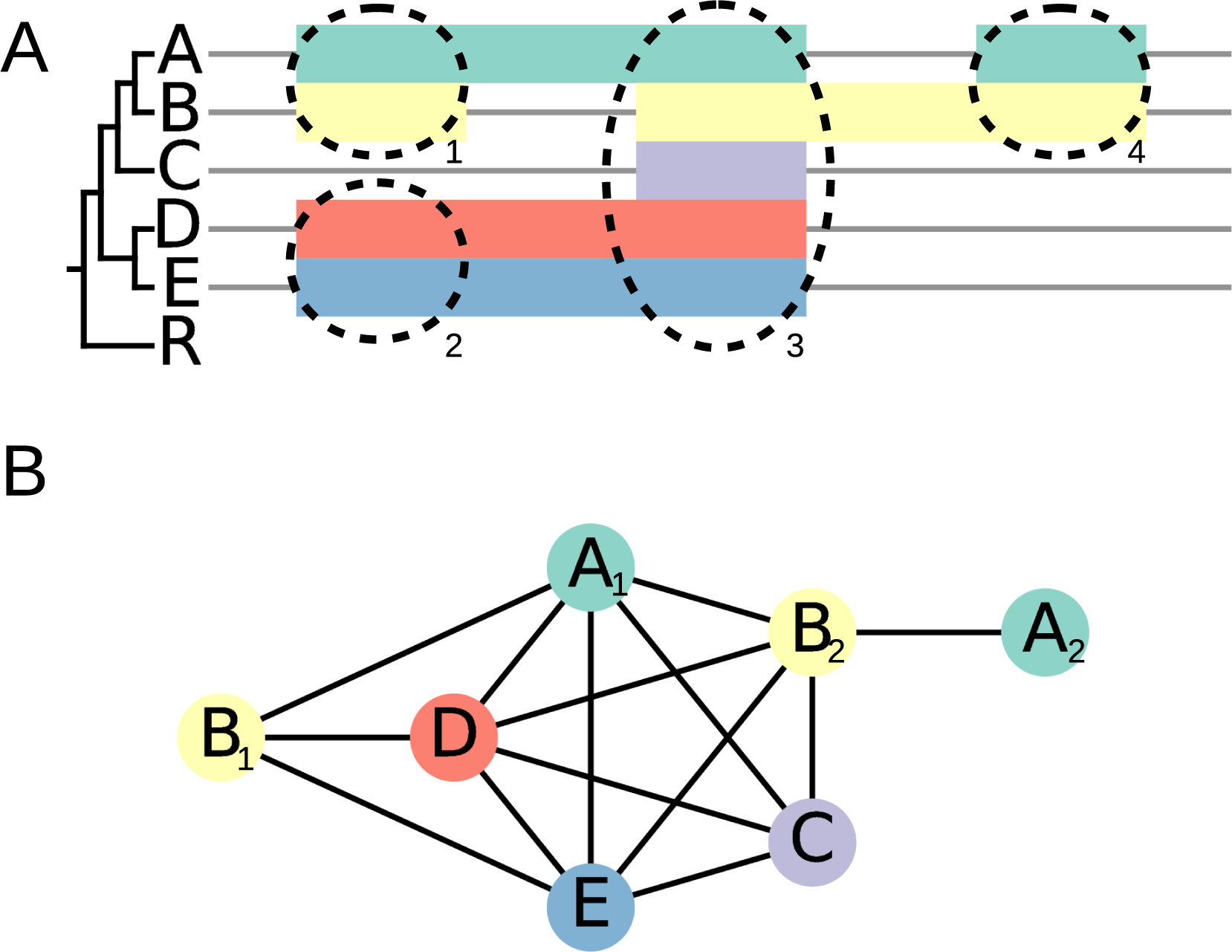
A) A toy alignment of 5 genomes (*A*, *B*, *C*, *D*, and *E*) to a reference genome (*R*). Each row (and color) corresponds to an alignment of a query to the reference. The horizontal position corresponds to the position in the reference genome. Grey lines represent orthologous segments between the reference and corresponding query, while colored bars represent qbreaks. The cladogram of the 6 genomes is shown on the left side, which is rooted on the branch leading to the reference. Dashed circles represent tbreaks {*τ*_1_*, τ*_2_*, τ*_3_*, τ*_4_} B) The interval graph of the qbreaks in A. Qbreaks are labeled and colored according to their query, and where more than one qbreak occurs in an alignment to a query they are indexed by their positions. Black edges are drawn between qbreaks that overlap each other. Note that tbreaks *τ*_3_ and *τ*_4_ are both maximal cliques (as can be seen in B) and maximal tbreaks (as can be seen in the tree on A), while *τ*_1_ is a maximal tbreak but not a maximal clique since {*A*_1_*, B*_1_*, D, E*} forms a clique. *τ*_2_ is neither a maximal clique nor a maximal tbreak, since it is a subset of *τ*_3_, but is discovered (and subsequently discarded) by our algorithm.

Finding the most parsimonious history of double-strand breaks for a given genome over the observed tree is equivalent to finding the smallest set of tbreaks that includes each qbreak at least once. We note that this solution may include a single qbreak in multiple tbreaks, corresponding to inferring multiple break events within the region of a qbreak. Such an inference is consistent with the data since qbreaks evidence at least one break within them. Multiple hits in the same region could be masked by a concomitant deletion associated with the region, or by sequence divergence following null functionalization at the site of the break. However, we note that this method does create a bias in the placement of double-strand breaks, that is, to make them more clustered.

#### 2.3.2 Synteny as an Interval Graph

Let the set of qbreaks be vertices in a graph where undirected edges represent an overlap between two qbreaks. Such a graph is known as an **interval graph**. We find it interesting to note that the queries of the qbreaks are a coloring of the interval graph, and the chromatic number of the graph is therefore at most the number of queries.

A tbreak is a set of fully connected qbreaks in the interval graph (i.e. a **clique**) whose members form a tree consistent partition. Finding the smallest set of tbreaks that includes each qbreak at least once is equivalent to finding a minimum cover using tbreaks. The minimum clique cover problem is NP-hard for arbitrary graphs, but can be solved in polynomial time for interval graphs. We follow a similar method to that described by Vandal *et al.* (1997) to enumerate all minimum covers of the interval graph using tbreaks.

A clique is considered **maximal** if it is not a proper subset of any other clique (i.e. it cannot be grown by adding another qbreak). A clique cover merely needs to contain each qbreak at least once, so can be composed of overlapping cliques. A small non-maximal clique in a cover can therefore always be replaced by a maximal clique that is its superset. A minimum clique cover composed of maximal cliques therefore exists. Minimum covers composed of non-maximal cliques can also exist, but the set of non-maximal cliques can be very large. In order to keep the problem tractable we consider only tbreaks that are maximal partitions of maximal cliques. This simplification leads to the breakpoint clustering bias previously mentioned and discussed at length later.

We refer to the multiple solutions to the tbreak cover problem as **histories**, as they represent multiple evolutionary histories that are equally parsimonious.

The unique feature of this problem relative to other minimum clique cover problems is the restriction of cliques to be tree consistent, rather than simply maximal. Once maximal cliques are found, they must be partitioned into their maximal tbreaks. Additionally, the method of Vandal *et al.* must be modified since maximal cliques have a strict ordering, while tbreaks are only partially ordered (Fig. 3). Fortunately, the low chromatic number of the qbreak interval graph ensures relatively low connectivity within the graph, leading to a single, obvious, best solution for large sections of the graph.

### 2.4 Estimation of the Break Rate

The resulting histories represent equally likely sets of inferred double stranded breaks that occurred within a known range of space (across the genome) and time (along the evolutionary tree). From these data, we seek to make a maximum likelihood estimate of the break rate across the genome. We compute such an estimate by calculating the joint likelihood across the four forms of uncertainty: 1) the underlying rate that produced the observed count of breaks, 2) the temporal placement of the break within each tbreak in the history, 3) the spatial placement of the break within each tbreak in the history, and 4) the selection of the history.

Due to the rarity of events, the resulting landscape is highly rugged, with high rates estimated at bonds where breaks are observed, and rates near zero estimated within unbroken regions. To make more even estimates of the break rate across a region, we apply a rectangular (sliding window) smoothing function to the counts of breaks across the genome and calculate likelihoods within the windows.

#### 2.4.1 Poisson Model of Double Strand Breaks

Starting with the uncertainty of the underlying rate, we model rearrangements as an independent Poisson process for each phosphodiester bond in the genome. Individual bonds break, or decay, at a constant rate, *λ*, and the break rates across the genome are denoted {*λ*_1_*, …, λ*_*N*_} where *N* is half the number of phosphodiester bonds in the genome. For a given region of the genome defined by the closed interval [*p, q*], the break rate within that region is estimated as the number of breaks observed, *c*, divided by the amount of evolutionary time that region is observed, *t*.

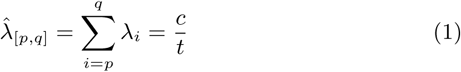

The maximum likelihood estimate of the break rate, *λ*_*i*_, for an individual bond within the region, *i ∈* [*p, q*], is equal to the total rate divided by the number of bonds.

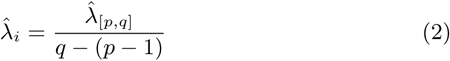

The likelihood function of the break rate for a region can be determined from the likelihood function for the mean number of breaks within the region. From the rate for the region, the mean number of breaks within the region is simply *µ*_[*p,q*]_ = *λ*_[*p,q*]_*t*. The likelihood function for the mean number of events in a Poisson process is

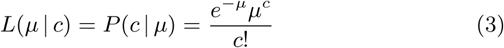

so it follows that:

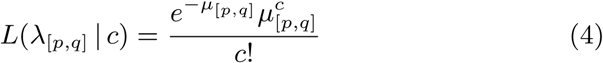

We use this model to compute the expectation and likelihood land-scape of *λ* for sliding windows across the reference genome.

#### 2.4.2 Observed Evolutionary Time

The time for which we observe a phosphodiester bond is equal to the combined length of branches of the phylogeny that can be inferred to share that bond. As a decay process, we cannot observe a bond after it is broken. However, we can still observe multiple breaks for a given bond, since speciation acts as a birth process for the bond (Fig. 4). We represent the observed evolutionary time for each bond in the genome as {*t*_1_*, …, t*_*N*_}, where *t*_*i*_ is dependent on the pattern of tbreaks at that bond.

**Figure 4:**
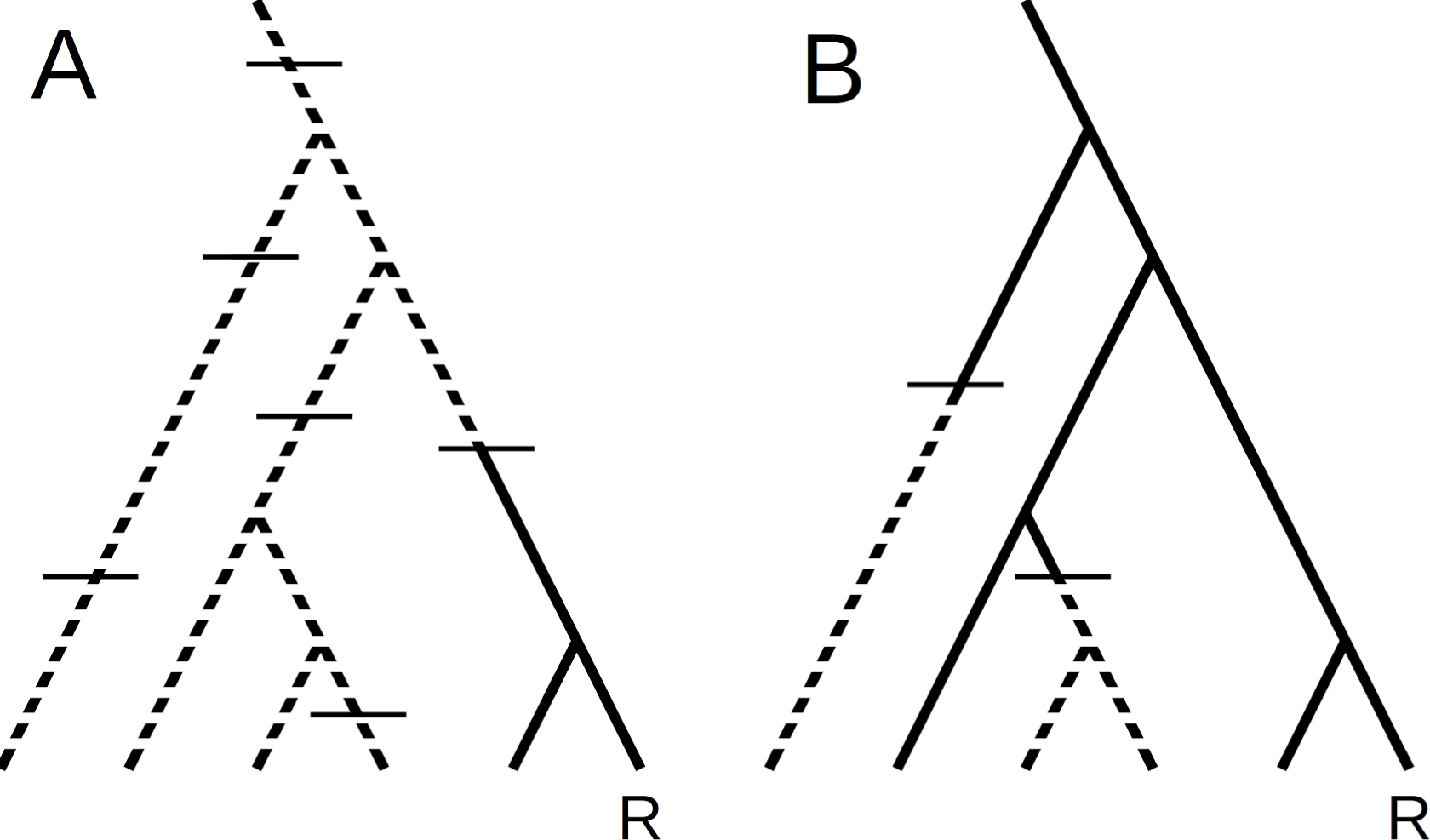
A hypothetical phylogeny showing break events at a rapidly breaking bond (A) and a relatively conserved bond (B). From the perspective of the reference state (labeled “R”), multiple hits beyond the most recent cannot be observed. The unobservable evolutionary time is masked (dashed lines) beyond these most recent events. The observed lifetime of the bond is equal to the sum of the lengths of the solid branches. At rapidly breaking bonds (A), a break in a recent ancestor of the reference is more likely to have occurred, and deeply branching queries provide no new information. At conserved bonds (B), however, deeply branching queries add significantly to the length of the tree observed and thus provide a greater chance for observing rare events.

The position of a break along the branch of a tbreak cannot be known, a condition known as “interval-censoring”. Assuming that new bonds after a break are exaclty as stable as the ancestral bonds or that breaks are rare events, the probability density function of the time to an event over an interval of time given that exactly one event occurred is approximately uniform. We therefore account for the temporal uncertainty in determining the evolutionary time observed for the region of the tbreak by simply using the midpoint of the tbreak’s branch, which is the expectation for the observed time.

When multiple tbreaks overlap spatially, each break masks the evolutionary history on the tree beyond the break from the perspective of the reference. For the region of overlap, we therefore calculate the total length of the masked branches and subtract it from the total length of the tree to determine the observed evolutionary time.

#### 2.4.3 Spatial Placement of Breaks within Tbreaks

Just as breaks cannot be precisely placed in evolutionary time, breaks are also interval-censored spatially. When two orthologous segments of a query are immediately adjacent to each other, the inferred qbreak between them consists of a single phosphodiester bond and the break can be precisely placed. Where there is a gap in the alignment, however, the placement of the break on any particular bond can not be known. Such qbreaks represent “hidden” breakpoints (Sankoff and Blanchette, 1998). This situation can arise in repetitive or other regions where our ability to align the genomes is poor, when the rearrangement is associated with an insertion or deletion, or where the sequence has degenerated around the site of the break following rearrangement. We expect degeneration of conserved elements at the break point of a rearrangement to be common since rearrangements often result in null-functionalization.

As in the larger genome, we expect the likelihood of a break to be heterogeneous across the bonds within a tbreak. In the absence of an adequate predictive model that incorporates both structural predilection and selective constraints, though, we use a uniform model for the spatial placement of breaks within a tbreak. The likelihood that a tbreak (*τ*) is an inference of a break within the interval [*p, q*] is simply the number of bonds that the interval overlaps with the tbreak divided by the number of bonds the tbreak covers. Or formally:

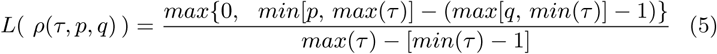

Given multiple tbreaks that overlap the region [*p, q*], denoted T_[*p,q*]_, the combinations of tbreaks that could be placed within the region is the powerset of T_[*p,q*]_. Let *χ ∈ 𝒫* (T_[*p,q*]_) denote a combination of tbreaks potentially in the region [*p, q*] and *n* be the number of overlapping tbreaks, *|*T_[*p,q*]_*|*. The joint likelihood for each possible combination of tbreak placements is the product of the likelihoods that each tbreak in the combination, *τ ∈ χ*, is placed in the region, *L*(*ρ*(*τ, p, q*)), times the product of the likelihoods that each tbreak not in the combination, *τ ∈* T_[*p,q*]_ *\ χ*, is not placed in the region, (1 - *L*(*ρ*(*τ, p, q*))), divided by the number of combinations, 2^*n*^:

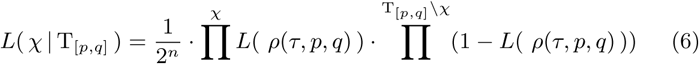

#### 2.4.4 Combining Histories

Given the set of *k* maximally parsimonious histories, *H* = *{h*_1_*, …, h*_*k*_*}*, the marginal likelihood of a given rate at a site is sum of the likelihoods over all histories. Under the paradigm of parsimony, though, we assume that all histories are equally likely, so:

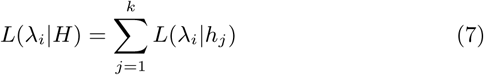

In computing the histories, though, it is easiest to compute a set of solutions for each disconnected (and therefore independent) subgraph of the qbreak graph. As described later, some tbreaks are obviously essential to all minimum cover solutions and their members can be effectively eliminated from the qbreak graph. The elimination of qbreaks from the graph can create disconnected subgraphs that may be solved independently, but whose qbreaks overlap spatially and which must be combined to achieve the full solution.

The resulting histories from the algorithm described below consist of a multiset of independent solutions, each of which is a multiset of minimum covers for a disconnected subgraph. The complete number of histories, being the product of the number of covers of each subgraph, has the potential to be prohibitively large. However, in estimating the break rate at a particular bond in the genome, those subgraphs that don’t include a qbreak that overlaps the bond can be disregarded, reducing to one or a few the number of subgraphs that need to be considered at a time.

For a qbreak graph that is composed of *κ* disconnected subgraphs, let Γ = {*γ*_1_*, …, γ*_*κ*_} be the family of minimum cover solutions for each disconnected subgraph of the qbreak graph. Each subgraph solution, *γ*, is itself the family of the minimum tbreak covers of the corresponding subgraph, and these covers are themselves sets of tbreaks. The set of histories is therefore equal to the unions of the *κ*-tuples of covers resulting from the *κ*-fold Cartesian product of all *γ ∈* Γ. Let *f* be this function such that *f* (Γ) = *H*.

For each region defined by a unique pattern of tbreaks [*p, q*], we remove all tbreaks that don’t overlap the region to form Γ_[*p,q*]_ which is the multiset of multisets of covers consisting only of tbreaks that overlap the region. We define the function, *g* such that *g*(Γ*, p, q*) = Γ_[*p,q*]_. Clearly, Γ_[*p,q*]_ should be equivalent to Γ within the region [*p, q*], so that*L*(*λ*_*i*_*|*Γ) = *L*(*λ*_*i*_*|*Γ_[*p,q*]_) where *i ∈* [*p, q*]. If we define *g* as a recursive function that removes all non-overlapping tbreaks from a multiset and any multiset within that multiset, then *g*(*f* (Γ)*, p, q*) = *f* (*g*(Γ*, p, q*)) = *H*_[*p,q*]_. Since *g* will completely empty most covers within the subgraphs, most *γ ∈* Γ_[*p,q*]_ will be a multiset of one or more empty sets. Any such subgraph, *γ*_*Ø*_, will be the multiplicative identity with regards to *L*(*λ*_*i*_*|f* (Γ_[*p,q*]_)), since *f* (*{γ*_*Ø*_*, γ*_*j*_*}*) will equal the multiset composed of *γ*_*j*_ unioned to itself *|γ*_*Ø*_*|* times. Remembering that all histories are assumed equally likely, this means *L*(*λ*_*i*_*|{γ*_*Ø*_*, γ*_*j*_*}*) = *L*(*λ*_*i*_*|{γ*_*j*_*}*), and all such subgraphs can be discarded in calculating *H*_[*p,q*]_.

#### 2.4.5 Consolidation of Uncertainty

We note that *c*, the number of breaks counted in the region [*p, q*], is equal to *|χ|*, the number of tbreaks placed in the region. Additionally, each history, *h ∈ H*_[*p,q*]_, is a possible set of tbreaks that could occur within the region [*p, q*], that is, a possible instance of T_[*p,q*]_. Summing across possible histories, *h*_*j*_ *∈ H*_[*p,q*]_, and again across the possible placement combinations, *χ*_*i*_ *∈ 𝒫* (*h*_*j*_), we combine equations 4, 6 and 7, above, to compute the likelihood of a given break rate, *λ* over the region [*p, q*]:

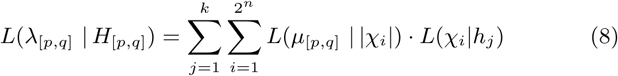

While the calculation of joint likelihoods of all combinations within the region takes *O*(*k*2^*n*^) time, *k* = *|H*_[*p,q*]_*|* and *n* = *|h*_*j*_ *∈ H*_[*p,q*]_*|* should be both be quite small except for large trees in regions of the genome that were ancestrally conserved but have diverged in multiple more recent lineages. This computational burden is multiplied by the sampling resolution: the number of rates for which the likelihood is computed, and the number of windows across the genome.

## 3 Algorithm to Enumerate Maximum Parsimony Histories

Here, we describe a method to solve the tbreak cover problem and enumerate all maximum parsimony histories. First, we recognize that all maximal tbreaks can be formed by partitioning the maximal cliques in the qbreak interval graph. After finding this set of plausible tbreaks, we reduce the size of the problem substantially in the manner of Vandal *et al.*, and finally recursively enumerate the minimum covers using these tbreaks.

### 3.1 Finding Maximal Cliques

Via transitivity, any tbreak that is an improper subset of a non-maximal clique must be contained in a maximal clique. Therefore we need only consider the maximal cliques of the qbreak interval graph. Finding all maximal cliques in a graph is known as the **clique problem**, which is well known to be NP-complete for arbitrary graphs, but is equivalent to a sorting problem for interval graphs. We use an algorithm very similar to the one described by Gupta *et al.* (1982), which follows:

First, we sort the left and right endpoints of all qbreaks together, with right endpoints preceding left endpoints when their positions are equal.

#### Algorithm 1

Exhaustive enumeration of minimum covers of an interval graph of ⊂ -minimal qbreaks. The choice of qbreak for *x* is arbitrary, as other qbreaks will be selected in the deeper levels of recursion. *x*^*\**^ is the set of maximal tbreaks *x* is affiliated with. The algorithm is initialized with a set of connected non-simplicial qbreaks, the simplicial qbreaks connected to the first set, and the essential tbreaks that cover the simplicial qbreaks.

~~~
minimum_ covers (qbreaks, covered_ qbreaks, accepted_tbreaks) :
         **if** qbreaks = : Ø
               **return** {accepted_tbreaks}
         **else**:
              covers ← Ø
              *x* ← **any** qbreak within qbreaks
              **for** *τ* **in** *x*^*^ :
                     remaining _qbreaks ← qbreaks \*τ*
                     new _covered ← co ve red_ qb rea ks ⋃ *τ*
                     new _accepted ← accepted_tbreaks ⋃ {*τ*}
                     new_ covers ← minimum_ covers (remaining_ qbreaks, new_ covered, new _accepted)
                     covers ← covers ⋃ new_ covers
              **return** all_ minimum (covers)
all _ minimum (covers) :
         *l* ← min{*|C|* : *C* ∈ covers}
         **return** {*C ∈* covers : *|C|* = *l*}
~~~

We proceed through the sorted list of endpoints, adding each qbreak to a set when we encounter its left endpoint, and removing it from the set when we encounter its right endpoint. At each left endpoint that immediately precedes a right endpoint we record the set as a maximal clique.

### 3.2 Partitioning Maximal Cliques into Tbreaks

We partition maximal cliques into tbreaks by finding their tree-consistent subsets.

Remembering that cliques are defined as fully connected sets of graph elements, we note that every subset of a clique is itself a clique. Furthermore, since the tree is hierarchical, for any two non-identical tbreaks that contain the same qbreak one must be a proper subset of the other, and therefore is not maximal. There is therefore a single partitioning of a maximal clique into maximal tbreaks which is equivalent to the minimum partition. With respect to the graph of qbreaks as a whole, however, some of these tbreaks may be non-maximal. These non-maximal tbreaks arise when neighboring maximal cliques are partitioned such that one tbreak is a subset of a neighboring tbreak. For example, in Fig. 3, *τ*_2_ is maximal within the clique {*A*_1_*, B*_1_*, D, E*} but not within the clique {*A*_1_*, B*_2_*, C, D, E*}. Such non-maximal tbreaks can be discarded following the same logic as for discarding non-maximal cliques.

To partion each maximal clique into its maximal tbreaks, we first find the origin of the reference state on the rooted tree by climbing from the reference leaf until we reach the ancestral branch that subtends all taxa with the reference state (i.e. that are not represented in the clique). If the origin branch is not the root of the tree, we place a tbreak on the origin branch and remove qbreaks with queries on the other side of the break from the clique. While qbreaks remain in the clique, we select a qbreak arbitrarily and climb the tree to find the highest branch that subtends only queries represented in the clique. We place a tbreak on that branch and remove all qbreaks with queries that are under it.

This process of partitioning maximal cliques into tbreaks can yield multiple instances of the same tbreak (i.e. tbreaks whose members are identical) in adjacent cliques. We cull these redundant tbreaks to produce the set of maximal tbreaks.

### 3.3 Enumerating Histories

From the set of plausible tbreaks, we enumerate all minimum covers in the manner of Vandal *et al.*. Prior to enumeration, Vandal *et al.* make several observations that enable a significant reduction of the size of the problem, which we summarize here.

First, we consider only covers of the ⊂**-minimal** qbreaks. We refer to the set of tbreaks that a qbreak belongs to as the **affiliations** of the qbreak, denoted *x*^*\**^, where *x* is a qbreak. Qbreaks whose affiliations contains no other set of affiliations as a proper subset are termed “⊂- minimal”. For example, in Fig. 3, *A*_1_ and *B*_2_ are not ⊂-minimal because their affiliations (*{τ*_1_*, τ*_3_*}* and *{τ*_3_*, τ*_4_*}*, respectively) are subset by the affiliations of *C* (*{τ*_3_*}*), which is ⊂-minimal.

The affiliations of a qbreak that is not ⊂-minimal contain exactly the affiliations of at least one ⊂-minimal qbreak. Each qbreak need only be covered by a single tbreak, so any tbreak that covers the ⊂-minimal qbreak also covers any non ⊂-minimal qbreaks whose affiliations contain the affiliations of the ⊂-minimal qbreak. Therefore, the minimum covers of the ⊂-minimal qbreaks are exactly the minimum covers of all the qbreaks. For example, in Fig. 3, any tbreak cover of *C* will also cover *A*_1_ and *B*2, so *A*_1_ and *B*_2_ can be ignored in finding minimum covers.

Second, we can include all essential tbreaks *a priori*. Qbreaks that are affiliated with a single tbreak are termed “**simplicial**”, and the tbreak that covers them is termed “essential”. All simplicial qbreaks are necessarily ⊂-minimal. We need simply append these essential tbreaks to covers of the non-simplicial qbreaks.

Third, we can find minimum covers of the disconnected subgraphs of the remaining non-simplicial, ⊂-minimal qbreaks and combine them to achieve the full solutions as described in section 2.4.4.

We then exhaustively enumerate the minimum covers for each disconnected sub-graph using algorithm 1, initialized with the set of qbreaks in the subgraph, the simplicial qbreaks adjacent to the graph, and the essential tbreaks, respectively. Vandal *et al.* first presented the substance of this algorithm as Algorithm 3.5, although with an error. We present the corrected form here, as well as modify the algorithm slightly due to the lack of a linear ordering on simplicial qbreaks (simplicial elements in an interval graph have a natural ordering) and to return only minimum, rather than all minimal, covers. For further details and proofs regarding the enumeration of minimal covers we refer the reader to Vandal *et al.*’s full paper.

## 4 Results

We implemented this algorithm and model in Python. BRAG depends upon the Python packages NumPy (Oliphant, 2006), Pandas (McKinney, 2010), StatsModels (Seabold and Perktold, 2010), SciPy (Jones *et al.*, 01), BioPython (Cock *et al.*, 2009), ETE (Huerta-Cepas *et al.*, 2016), and Matplotlib (Hunter, 2007).

We used BRAG to estimate the rates of double-strand breaks across the finished genome of *N. crassa* by comparing it to draft genomes of 14 *Neurospora* species and an outgroup, *Sordaria macrospora*. BRAG took only 6.5 min to run on a single thread of an Intel Xeon E5-2670 v2 CPU, and used 3.5 GB peak memory. However, solving the minimum clique cover by tbreaks (MCCT) problem took only 3 min 33 s and 494 MB peak memory, with the remaining resources used to draw figures.

We identified 15,450 true (19,544 true or false) qbreaks and 12,356 true (16,176 true or false) tbreaks. The tbreaks formed 10,801 true (14,409 true or false) disconnected subgraphs. Only 23 true (25 true or false) subgraphs had multiple maximally parsimonious solutions, resulting in 60 true (77 true or false) solutions to the MCCT problem. The resulting estimates of the break rate for chromosome 2 is shown in Fig. 5.

**Figure 5:**
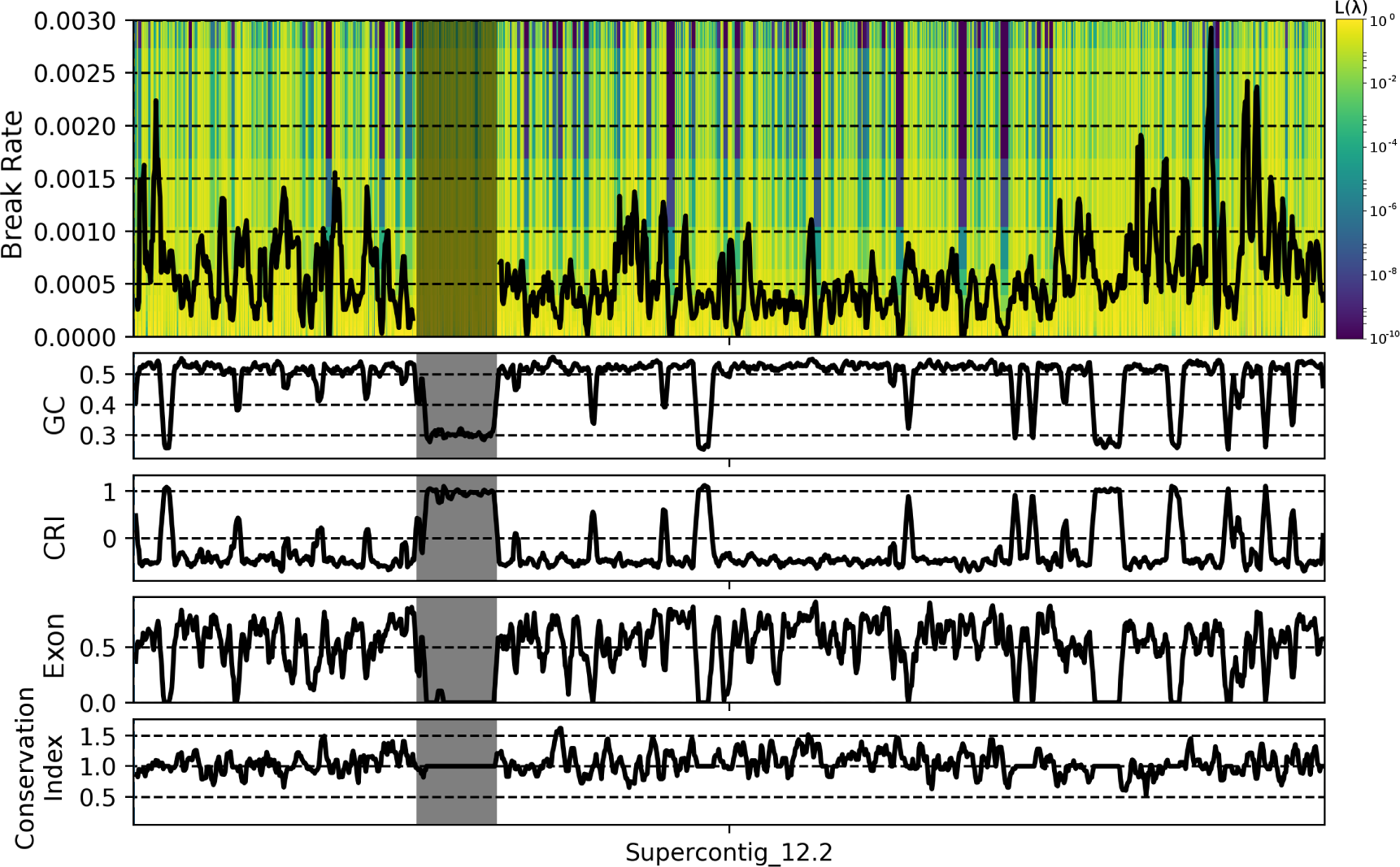
Likelihood landscape of break rates along chromosome 2 of *N. crassa*. Yellow indicates high likelihood, while purple indicates low likelihood. The black line shows the maximum likelihood estimate of the break rate for sliding windows with a step of 4,000bp and width of 20,000bp. The break rate was not estimated within the centromere, which is shaded on all tracks. Sliding window calculations of the G/C content, Composite RIP Index (a measure of repetitive content), portion of exonic sequence, and a conservation index are shown below the break rate. The conservation index is calculated as described in section 4.2

The maximum likelihood estimate of the break rate when only true qbreaks were used was identical to the estimate when true and false qbreaks were used for 12.4% of sliding windows. 89.8% of non-identical windows showed an elevated rate when using both true and false qbreaks, with a mean increase of 42.9%. The over-estimation of the break rate when using false qbreaks was stronger in regions with a lower break rate (*p <* 10^-126^, Fig. S1), indicating the higher ratio of poorly assembled regions to real qbreaks in these regions.

### 4.1 Factors Affecting the Break Rate

We predicted that breaks would be more common in regions with high repetitive content due to the presence of transposable elements, and less common in gene-dense regions. In *Neurospora*, RIP silences transposons by selectively mutating CpA dinucleotides to TpA dinucleotides in nonunique sequences (Cambareri *et al.*, 1989). As such, regions that have been repetitive in *Neurospora*’s past bear a genetic signature, summarized by the Composite RIP Index (CRI) (Lewis *et al.*, 2009). Positive CRI values indicate RIP action, and thus repetitive sequence, while zero or negative CRI values indicate unique sequence.

We performed multiple linear regression analysis to identify factors correlated with the break rate. While a model incorporating G/C content, CRI, the portion of protein coding sequence, the portion of exonic sequence, and an index of the phylogenetic conservation of genes within a region (described below) found all factors to be significant (*p <* 0.001), the combined effect only explained 5.4% of the variation. Due to the action of RIP, all factors are significantly collinear because RIP enriches the genome for GC content while nullifying genes. We therefore performed independent linear regression between the break rate and CRI, exon density for nonrepetitive regions with CRI *<* 0, and conservation index. While significant, the effects of both exon density and CRI were vanishingly slight (*R*^2^ = 0.009 and 0.0005, respectively). Only the level of phylogenetic conservation of the genes within a region had any appreciable effect on the break rate (*R*^2^ = 0.029 in the independent linear model).

We looked for evidence of rapidly evolving subtelomeric regions by performing linear regression between the distance to the telomere and the break rate for each of the 14 chromosomal arms. After Benjamini-Hochberg correction for a false-discovery rate of 0.05, we found a significant negative relationship between distance from the telomere and the break rate for all but one of the 14 chromosome arms (Fig. S2A). A 0.5-1 Mbp region of elevated breakage is apparent on the most distal tips of many of the chromosome arms (e.g. the left arms of chromosomes 1, 6, and 7, and the right arms of all chromosomes).

Overall, we find evidence that GC content, repetitive content, and gene density have weak effects, while the level of phylogenetic conservation of genes in a region and the position along the chromosome arm have larger effects. However, collinearity between factors obstructs inference of independent effects and may be leading to spurious observations in the minor factors. Repetitive content was also negatively correlated with distance to the telomeres in 9/14 chromosome arms, which may explain the slight correlation between CRI, GC content, and the break rate. Similarly, our conservation index was positively correlated with distance to the telomeres in 12/14 chromosome arms, indicating linkage between novel genes and rearrangements in a spatially organized manner (Fig. S2B).

### 4.2 Fragile Regions Are a Reservoir of Novel Genes

If rapidly rearranging (i.e. fragile) regions of the genome serve as engines for genetic novelty, then we hypothesized that fragile regions will harbor more species-specific genes that could be involved in recent adaptation while stable regions of the genome will harbor more highly conserved genes. To test this hypothesis, we categorized each gene in the genome by the phylogenetic level at which the gene is lineage specific, as reported by Kasuga *et al.* (2009). Kasuga *et al.* reported 5 levels of lineage specificity: species (*N. crassa*), subphylum (Pezizomycotina), phylum (Ascomycota), subkingdom (Dikarya), and all of cellular life.

We calculated a conservation index for regions in the genome as 1 minus the exon density of genes unique to *N. crassa* plus the exon density of genes common to all life. The resulting scale ranges from 0, indicating a region composed entirely of species-specific exons, to 2, indicating a region composed entirely of highly conserved exons. Values near 1 indicate either little exonic sequence or an even mix of species-specific and highly conserved exons.

We then ranked all the genes in *N. crassa* by the break rate of the window they occur in and compared the distribution of ranks for genes from each level of phylogenetic specificity (Fig. 6A). The mean break rate ranks, from species-specific to common in all cellular life, were 2,162, 2,502, 2,764, 2,386, and 2,614. We rejected the null hypothesis that the distribution of break rates was identical for each level of phylogenetic specificity with a Kruskal-Wallis test (*p <* 10^-24^), and species-specific genes had the fastest mean rank break rate.

**Figure 6:**
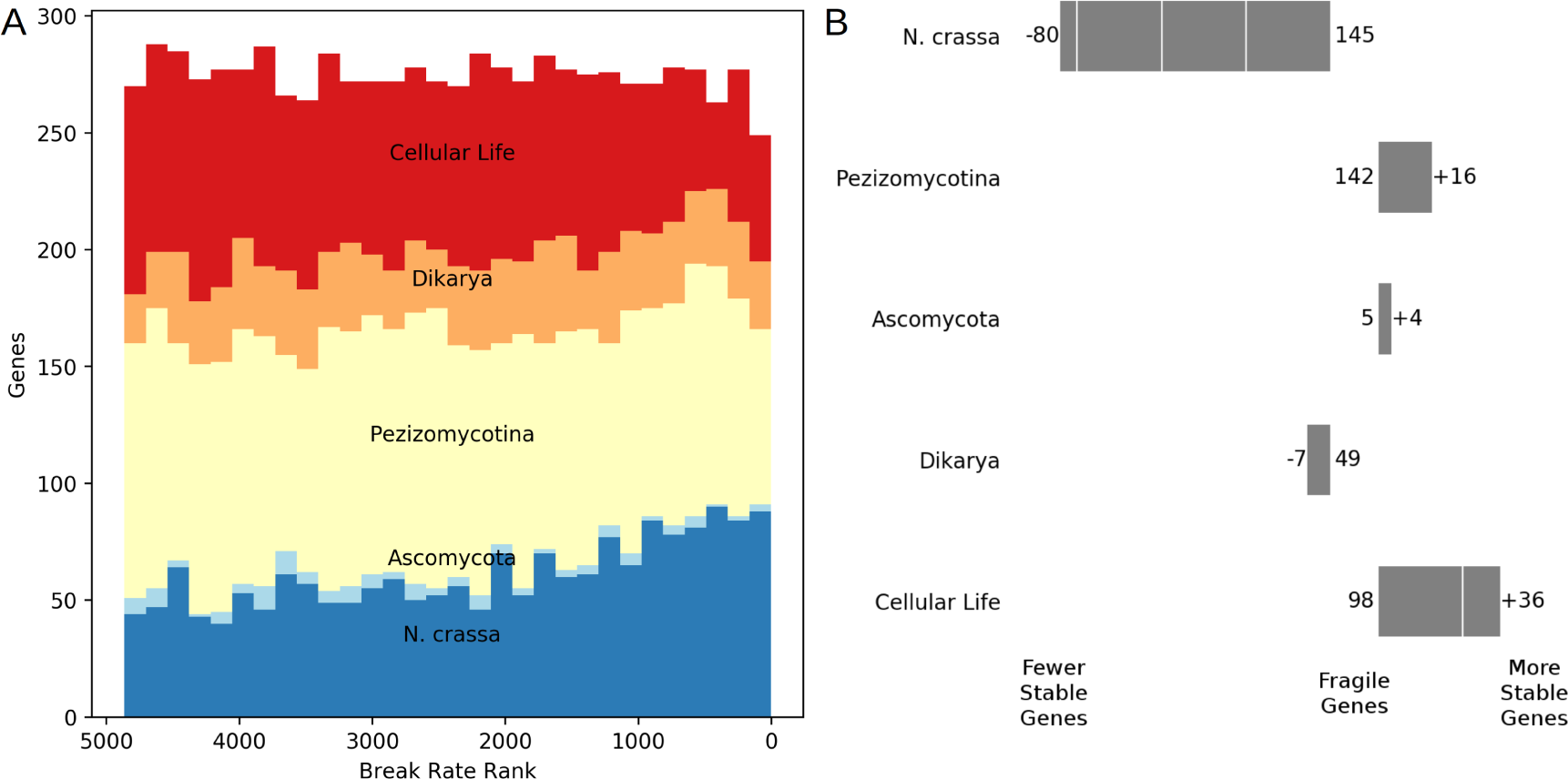
A) Stacked histograms of the number of genes within the five levels of phylogenetic specificity at different break rate percentiles show that the distribution of break rates around *N. crassa* specific genes is higher than more highly conserved genes. B) The difference in number of genes from each category is most apparent between the most fragile (right side in A) and most conserved (left side in A) regions of the genome. For each level of phylogenetic specificity, the difference between the number of genes in the most stable 2 Mbp of the genome and in the most fragile 2 Mbp is shown. The number of genes of each category in fragile regions is given in the center.

This pattern is especially apparent between the most fragile and most conserved regions of the genome. The most fragile 2 Mbp (about 5%) of the genome contained 553 genes, 451 of which were phylogenetically classified, and the most stable 2 Mbp of the genome contained 488 genes, 414 of which were phylogenetically classified. The most fragile regions had roughly the same number of genes as the most stable regions for genes that were specific at the subphylum, phylum, and subkingdom levels, but the fragile regions had far more species-specific genes while the stable regions had far more genes that were common to all cellular life (Fig. 6B).

### 4.3 The Two-Speed Genome of *Neurospora crassa*

Overall, we find that rearrangements abound across the genome of *N. crassa*, especially within the subtelomeric regions involved in recent adaptation. As has been described in humans (Linardopoulou *et al.*, 2005) and plasmodium (Cerón-Romero *et al.*, 2018), we observe subtelomeres that are enriched for rearrangements and recently evolved genes and are approximately 200 Kbp to 1 Mbp long.

Selection plays a clear role in filtering out rearrangements that occur within genes while permitting rearrangements in intergenic regions. The subtelomeres, though, show a propensity for rearrangement beyond this selective filter. If the elevated break rate within the subtelomeres were due merely to an increased tolerance for gene disruption then we would expect subtelomeres to be relatively gene sparse. To the contrary, we observe more genes in the most highly rearranged portions of the genome than in the most stable. It appears that selection not only filters out rearrangements that land within genes, but also favors a higher level structure of where rearrangement occurs.

Furthermore, the manner in which rearrangement creates novel genes for adaptation appears structured, since the placement of species-specific genes is extremely non-random (*χ*^2^ = 5922, df = 1, p = 0). Species-specific genes are far more likely to be surrounded by other species-specific genes than they are to neighbor a highly conserved gene. In comparing the neighborhoods of the 4,675 *N. crassa* specific genes and 6,967 genes common to cellular life, if the genome were randomly ordered we would expect 1,866 pairs of specific genes to be neighbors, 4,176 pairs of conserved genes to be neighbors, and 5,582 divergent pairs to be neighbors. In actuality, though, there are 3,858, 6,168, and 1,598 pairs of each category, respectively.

Still, the majority of variation in the break rate remains unexplained by the factors examined here, and this may reflect the heterogeneous effects of selection as well as biases in the patterns of incomplete data. However, the consistent partitioning of genes into rapidly evolving or conserved regions by organisms across Eukarya leads us to believe that there is significantly more to be learned about how organisms control the movement of genes throughout their genome.

Rearrangements can be an important source of adaptive variation (Brown *et al.*, 2010; Cerón-Romero *et al.*, 2018; Linardopoulou *et al.*, 2005; Miles *et al.*, 2016; Steenwyk and Rokas, 2018; Soucy *et al.*, 2015), and this source appears to be cultivated by eukaryotic genomes. In primates and likely other organisms, the structured repetition of subtelomeric regions facilitates rearrangements between rapidly evolving genes, while transposons aid in the transfer of adaptive genes in fungal pathogens. In *N. crassa*, an organism that keeps a meticulously clean house through mechanisms such as RIP, the permissibility of rearrangement in the subtelomeres suggests that they are gardens of variation, rather than wild landscapes.

## 5 Discussion

Our implementation of BRAG has demonstrated that it is possible to take a comprehensive look at the rearrangement landscape of a genome with relatively little computational power. By remaining agnostic about the mechanisms of rearrangements BRAG should scale highly favorably with the addition of more or more distantly related genomes. However, BRAG’s results are highly dependent upon correct inference of orthology and the underlying phylogenetic tree, and this scalability comes at the cost of potential idiosyncrasies in the results.

### 5.1 Scalability of BRAG

In their 2009 paper, Vandal *et al.* prove that the complexity of the algorithm to enumerate minimal covers is *O*(2^*m-*2^), where *m* = *|*T*|*, or the number of tbreaks in the graph. Solving disconnected subgraphs of T independently thus provides a significant advantage for this exponential time algorithm and allows for a roughly linear increase in computational time with increasing genome size. We further suspect that, for qbreak graphs, the rate at which *m* increases as new genomes are added will be slow because of the hierarchical nature of evolutionary trees.

Given the phylogeny relating the reference and queries, for any region of the reference genome the lowest nonzero maximum likelihood estimate of the break rate is equal to one divided by the product of the total tree length and the length of the region. This represents a kind of limit of detection for the break rate, since while the maximum likelihood estimate for an unbroken region is 0, from a biological perspective the break rate is more reasonably regarded as less than this limit. Much of the likelihood landscape across the genome is composed of regions without breaks, with a maximum likelihood of 0 and a negative slope determined by the width of the region. Improving the sensitivity of the break rate estimate is best achieved by adding genomes to the analysis that are distantly related to all other individuals, as deeper branches added to the tree add more evolutionary time to the tree overall, increasing the quotient of the limit of detection (Fig. 4).

At the same time, adding deeply branching genomes is not likely to add much complexity to the tbreak graph. The tbreak graph is, tautologically, sparse in conserved regions, so additional tbreaks are unlikely to tie Gordian knots. In more rapidly breaking regions deeply branching genomes are unlikely to provide information, since they are more likely fall behind an existing inferred tbreak and be effectively masked (Fig. 4). Adding deep-branching genomes thus provides more information about conserved regions of the genome by providing more opportunity to observe rare events, but does not provide much information about rapidly rearranging regions.

Within regions of the genome that have at least one qbreak, the accuracy of placing tbreaks on the tree is limited to the discrete internode distances (the branch lengths). Shorter internal branches allow for more precise placement of tbreaks, and thus more precise estimates of the break rate. The additional internally branching genomes are similarly unlikely to increase the complexity of the tbreak graph, but rather just to refine the placement of tbreaks.

### 5.2 Effects of Poorly Resolved Trees and Paralogy

While the inclusion of more data should generally yield better results, including ambiguously placed genomes in the analysis must be avoided. An incorrect tree topology will systematically duplicate tbreaks, leading to wild overestimates of the break rate. Such a situation can arise when there is recombination between individuals included in the analysis. Recombination results in different trees being true for different regions of the genome. Thus, a systematic bias will be introduced for regions that do not match the consensus tree. Including 3 individuals from a recombining population with architectural variation could nearly double the estimate of the break rate for half the genome. Horizontal transfer between species in the analysis could similarly multiply the break rate within the transfered region. Recombination will also lead to poorly resolved branches, which is why BRAG is only suitable for high-confidence phylogenies, or when regions that don’t match the consensus are masked (e.g. the centromeres in this analysis).

Similarly, paralogous segments that are misidentified as orthologous can lead to systematic overestimation of the break rate. Such a situation could occur following gene duplication and pseudogenization. If the orthologous copy is pseudogenized, then alignment may identify the working paralogue as orthologous, leading to three breaks observed where there should be one. Alignment methodologies that either allow for multiple homology relationships (e.g. cactus graphs) or that favor synteny over sequence identity should therefore be used. The former solution is particularly appealing. Although reading cactus alignments is not implemented by BRAG, we believe a method that traces paths on cactus graphs to identify tbreaks, combined with BRAG’s tbreak placement on trees and Poisson rate modeling would be particularly promising.

### 5.3 Effects of Genome Assembly Quality

BRAG is sensitive to the quality of the genome assemblies used, although we think the substantial similarity in break patterns using only “true” or both “true” and “false” qbreaks indicates that the general trends are robust to this sensitivity. In simulations using a similar framework, Zheng and Sankoff (2016) found detection of rearrangement events to be surprisingly robust to assembly fragmentation. Still, while we use estimates using only “true” or both “true” and “false” qbreaks as lower and upper bounds, respectively, this characterization is not entirely accurate since the presence or absence of a qbreak can complete, disrupt, grow, or shrink a tbreak. Zheng and Sankoff describe a superior treatment of ambiguous breaks: converting it into greater temporal censoring by allowing tbreaks to be placed across a subset of valid branches.

### 5.4 Limitations and Biases

BRAG remains agnostic about the mechanism and nature of the underlying rearrangements by measuring the age of existing bonds, with the assumption that younger bonds replaced more fragile bonds in the past. In balanced rearrangements, where two bonds are broken and the ends rejoined in a new arrangement, this assumption makes intuitive sense. But in unbalanced rearrangements where the total number of bonds is increased or decreased the number of tbreaks observed does not necessarily match the number of double strand breaks or joins that occurred. Insertions break one bond and rejoin the ends with a series of new bonds, but are observed as a single large tbreak. Conversely, deletions break two bonds and join the ends in a single bond, but are still observed as a single tbreak, this time of size 1. The point estimate of the break rate at the site of the deletion will be much higher than the insertion due to the tbreak’s smaller size, despite the fact that the events are reversals of each other. However, the estimates should be substantially similar after smoothing (e.g by sliding windows), and we believe that BRAG captures the essential nature of fragility as the evolutionary tolerance of a region for rearrangement.

BRAG favors choosing tbreaks that are higher up on the tree (i.e. closer to the reference) over tbreaks that are closer to the queries when both choices are equally parsimonious. This preference is due to only considering tbreaks that are maximal partitions of maximal cliques in our search for covers. Such a situation arises in Fig. 3. *τ*_1_ could be equally well explained by a tbreak on the ancestral branch of A and B or by a tbreak on the branch leading to B. Our algorithm dismisses the latter tbreak and favors the former tbreak because is is maximal.

This limitation results in two biases: an overestimation of the break rate in such regions, and a clustering of inferred breakpoints, such that the overestimation is much greater in one region while the neighboring region is underestimated. The overestimation of the break rate is due to selecting the highest possible placement of the tbreaks, and thus inferring the shortest possible observed evolutionary time in that region. The clustering bias arises because the placement of the tbreaks is bounded by the overlapping region of its qbreaks, so the two tbreaks evidenced by the shared qbreak will tend to be closer together. For example, in Fig. 3, if *min*(*A*_1_) *> min*(*B*_1_) then the choice of placement for the tbreak would lie between *min*(*A*_1_) and *max*(*B*_1_), while if *min*(*A*_1_) *≤ min*(*B*_1_) then the choice of placement for the tbreak would be the same as a tbreak on the branch leading to B (i.e. between *min*(*B*_1_) and *max*(*B*_1_)).

There are biological reasons to favor a clustering bias. Indeed, we add to the evidence that rearrangements are more common in domains involved in recent adaptation. However, this bias would be better modeled explicitly. We acknowledge this limitation to our methodology, but hope that BRAG will provide better resolution on the placement of break sites, enabling future models to account for these and yet unsupposed effects. With additional work, it may be possible to explore histories utilizing these non-maximal tbreaks.

### 5.5 Assumptions of BRAG

In addition to exploring non-maximal tbreaks, it may be advantageous to explore non-minimum histories. The restriction to minimum histories is imposed by the assumption of maximum parsimony, which is itself a weak assumption. A better method would employ a purely maximum likelihood approach, integrating across both minimum and less likely histories. Such a set of solutions could be very large, and a sampling methodology like that described by Vandal *et al.* could be utilized to explore the space of histories in a manner similar to Bayesian tree exploration and produce a more complete probabilistic model of the evolutionary history.

In addition to assuming a maximum parsimony history, BRAG assumes that the reference genome sequence and the organismal ecology remain unchanged throughout it’s evolutionary history. In reality, the evolving sequence of the genome (as well as the structure itself) influence the rearrangement dynamics. Rearrangements may become common in a region along a lineage following null-functionalization or an alteration of the epigenetic structure. Our inference of the rearrangement rate in such a region does not reflect the evolutionary dynamics of the reference itself, but rather an integration of it’s historical and historical-adjacent states that are found in the tree.

Similarly, demographic changes, and external or internal ecological shifts across the tree can change the rearrangement dynamics. Such a shift is seen in *Neurospora* where species have consistently but independently transitioned from self-sterility to self-fertility (Nygren *et al.*, 2011). This shift is associated with profound changes in evolutionary paradigm (Gioti *et al.*, 2013; Hann-Soden *et al.*, prep). As such, our results in *N. crassa* should be seen as a partial integration of the effect of self-fertility, induced by this lineage’s proclivity for transitions to self-fertility.

The ability to assay rearrangement dynamics on a large scale, facilitated by advances in sequencing and analysis methodologies, brings new questions into the fields of molecular biology, evolution, and ecology, such as the effect of sexual transitions in *Neurospora*. In particular, we believe there is still much to learn about evolution within the fields of molecular and ecological phylogenomics.

## 6 Data Access

BRAG, and all code used in the analysis presented in this paper, are available at https://github.com/channsoden/BRAG. All code is licensed under the 2-clause BSD license. Genomes and alignments used in this study are available at https://osf.io/ak54t/. Sequence reads for genomes sequenced by us are available from the NCBI (ACCESSION NUMBER GOES HERE).

Our pipeline for the whole-genome phylogenetic analysis can be found at https://github.com/channsoden/hannsoden-bioinformatics/WholeGenomePhylogeny and our rewrite of TIGER can be found at https://github.com/channsoden/hannsoden-bioinformatics/TIGER.

All strains used in this paper are available from the Fungal Genetics Stock Center at Kansas State University, Manhattan, Kansas.

## 7 Acknowledgements

This work was funded by NSF grant DEB-1257528, and CHS received support from the Philomathia Graduate Student Fellowship in the Environmental Sciences. This work used the Vincent J. Coates Genomics Sequencing Laboratory at UC Berkeley, supported by NIH S10 OD018174 Instrumentation Grant. Computation was performed on Berkeley Research Computing and Lawrence Berkeley National Lab’s HPC cluster accessed through the Computational Genomics Research Laboratory.

## Supplementary Information

**Figure S1:**
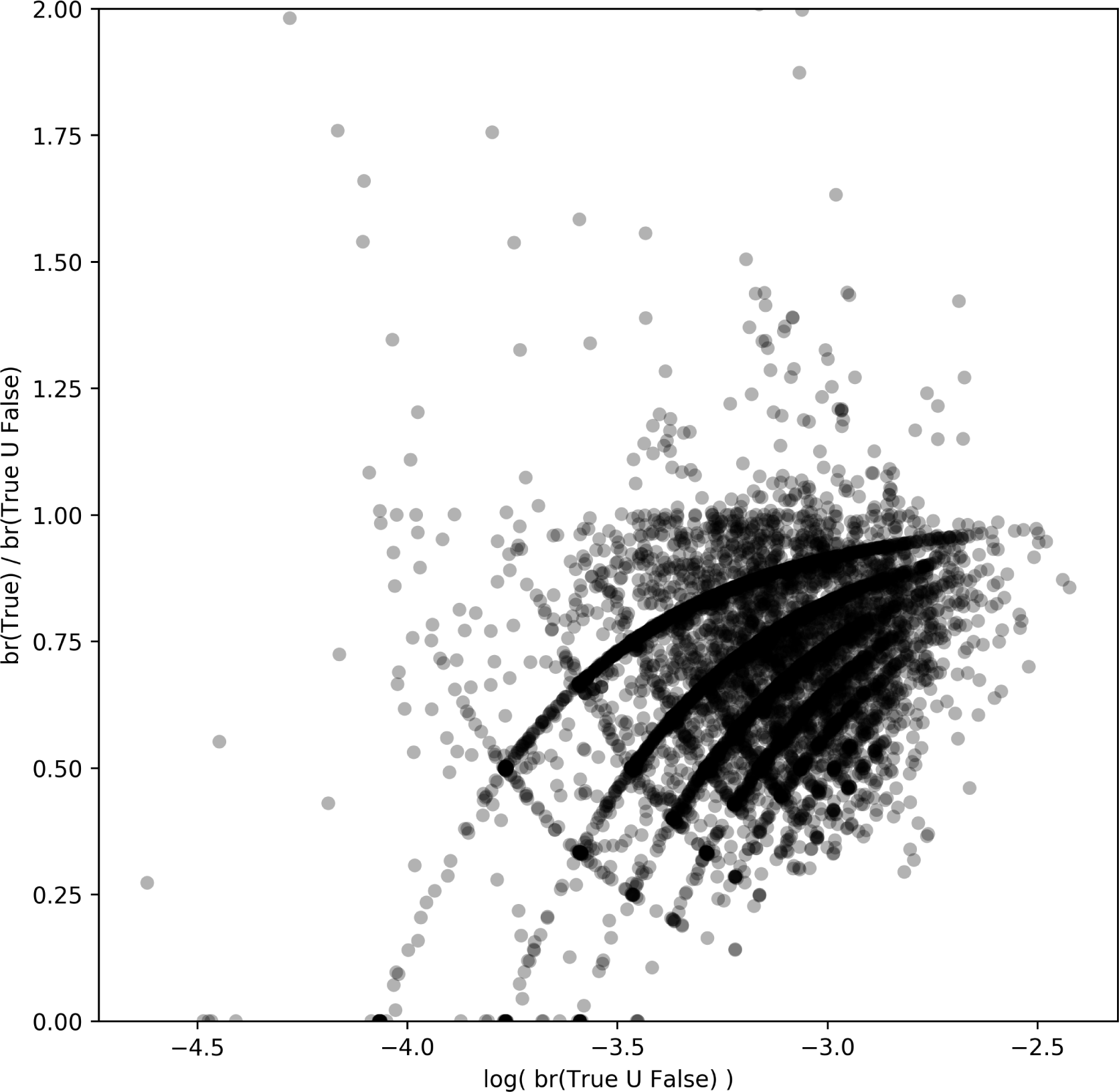
Using false qbreaks leads to greater overestimation of the break rate in regions with a low break rate. Each point represents the maximum likelihood estimate of the break rate across a 20,000bp window. Values greater than 1 represent windows where including false qbreaks led to a lower estimation of the break rate, while values lower than 1 represent windows where including false qbreaks led to a greater estimate of the break rate. The 12.4% of windows where the estimate did not depend upon false breaks are not shown.

**Figure S2:**
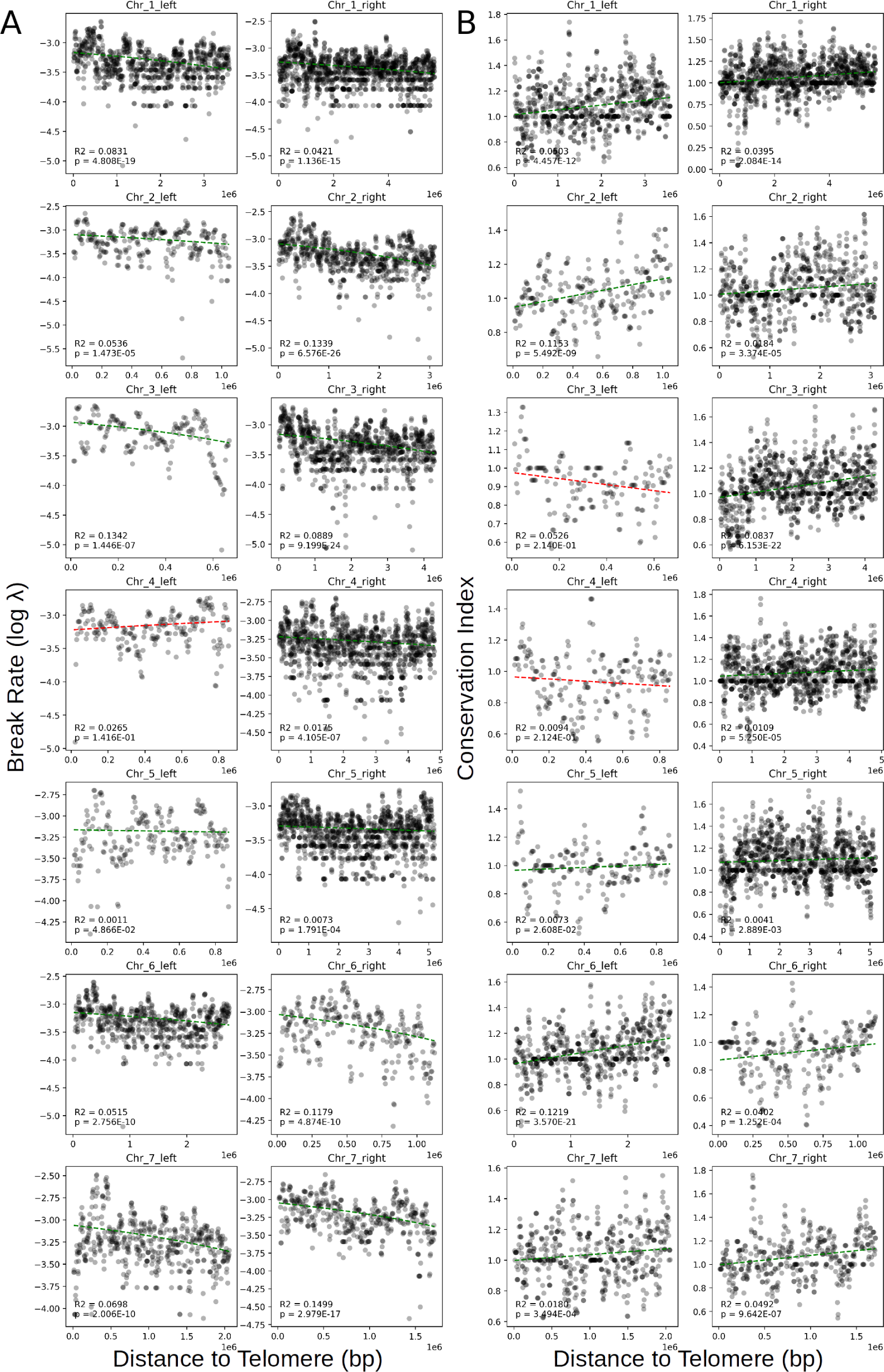
A) Break rate estimates (*λ*) are higher in the distal regions of chromosomes and lower toward the centromeres. B) The distal regions of chromosomes are enriched for species-specific genes, while the proximal regions are enriched for core conserved genes. Linear models are shown as green dashed lines when significantly negative, and red dashed lines when not. Distances are in millions of base pairs.

**Table S1:**
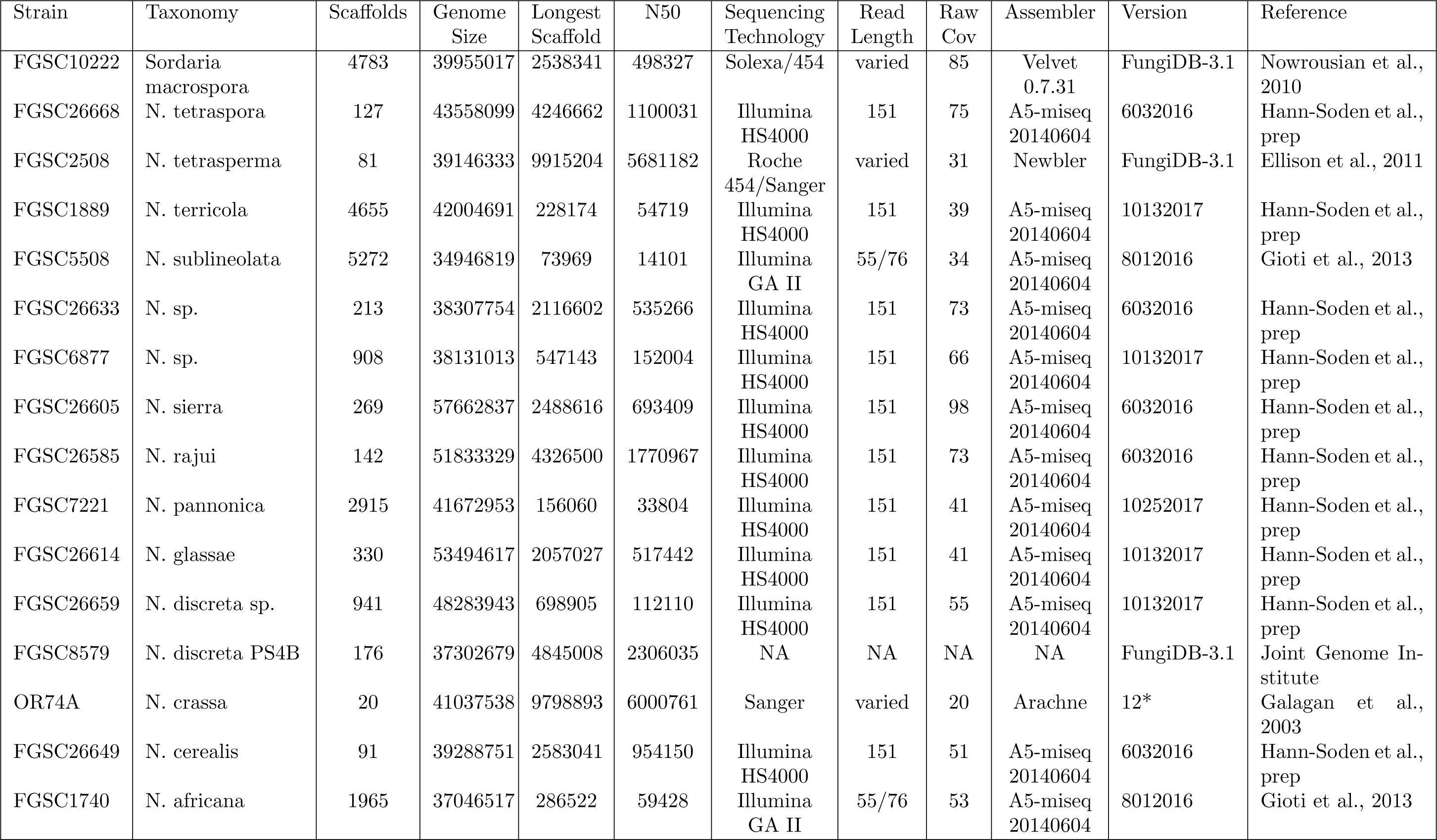
Genome assemblies used in this paper.

| Strain | Taxonomy | Scaffolds | Genome Size | Longest Scaffold | N50 | Sequencing Technology | Read Length | Raw Cov | Assembler | Version | Reference |
| --- | --- | --- | --- | --- | --- | --- | --- | --- | --- | --- | --- |
| FGSC10222 | *Sordaria macrospora* | 4783 | 39955017 | 2538341 | 498327 | Solexa/454 | varied | 85 | Velvet 0.7.31 | FungiDB-3.1 | Nowrousian et al., 2010 |
| FGSC26668 | *N. tetraspora* | 127 | 43558099 | 4246662 | 1100031 | Illumina HS4000 | 151 | 75 | A5-miseq 20140604 | 6032016 | Hann-Soden et al., prep |
| FGSC2508 | *N. tetrasperma* | 81 | 39146333 | 9915204 | 5681182 | Roche 454/Sanger | varied | 31 | Newbler | FungiDB-3.1 | Ellison et al., 2011 |
| FGSC1889 | *N. terricola* | 4655 | 42004691 | 228174 | 54719 | Illumina HS4000 | 151 | 39 | A5-miseq 20140604 | 10132017 | Hann-Soden et al., prep |
| FGSC5508 | *N. sublineolata* | 5272 | 34946819 | 73969 | 14101 | Illumina GA II | 55/76 | 34 | A5-miseq 20140604 | 8012016 | Gioti et al., 2013 |
| FGSC26633 | *N. sp.* | 213 | 38307754 | 2116602 | 535266 | Illumina HS4000 | 151 | 73 | A5-miseq 20140604 | 6032016 | Hann-Soden et al., prep |
| FGSC6877 | *N. sp.* | 908 | 38131013 | 547143 | 152004 | Illumina HS4000 | 151 | 66 | A5-miseq 20140604 | 10132017 | Hann-Soden et al., prep |
| FGSC26605 | *N. sierra* | 269 | 57662837 | 2488616 | 693409 | Illumina HS4000 | 151 | 98 | A5-miseq 20140604 | 6032016 | Hann-Soden et al., prep |
| FGSC26585 | *N. rajui* | 142 | 51833329 | 4326500 | 1770967 | Illumina HS4000 | 151 | 73 | A5-miseq 20140604 | 6032016 | Hann-Soden et al., prep |
| FGSC7221 | *N. pannonica* | 2915 | 41672953 | 156060 | 33804 | Illumina HS4000 | 151 | 41 | A5-miseq 20140604 | 10252017 | Hann-Soden et al., prep |
| FGSC26614 | *N. glassae* | 330 | 53494617 | 2057027 | 517442 | Illumina HS4000 | 151 | 41 | A5-miseq 20140604 | 10132017 | Hann-Soden et al., prep |
| FGSC26659 | *N. discreta* sp. | 941 | 48283943 | 698905 | 112110 | Illumina HS4000 | 151 | 55 | A5-miseq 20140604 | 10132017 | Hann-Soden et al., prep |
| FGSC8579 | *N. discreta* PS4B | 176 | 37302679 | 4845008 | 2306035 | NA | NA | NA | NA | FungiDB-3.1 | Joint Genome Institute |
| OR74A | *N. crassa* | 20 | 41037538 | 9798893 | 6000761 | Sanger | varied | 20 | Arachne | 12* | Galagan et al., 2003 |
| FGSC26649 | *N. cerealis* | 91 | 39288751 | 2583041 | 954150 | Illumina HS4000 | 151 | 51 | A5-miseq 20140604 | 6032016 | Hann-Soden et al., prep |
| FGSC1740 | *N. africana* | 1965 | 37046517 | 286522 | 59428 | Illumina GA II | 55/76 | 53 | A5-miseq 20140604 | 8012016 | Gioti et al., 2013 |

